# Spatiotemporal variation in drivers of parasitism in a wild wood mouse population

**DOI:** 10.1101/2020.05.16.099481

**Authors:** A.R. Sweeny, G.F. Albery, S.V. Venkatesan, A. Fenton, A.B. Pedersen

**Affiliations:** Institute of Evolutionary Biology, School of Biological Sciences, University of Edinburgh, Edinburgh UK; Department of Biology, Georgetown University, Washington, DC; Institute of Integrative Biology, University of Liverpool, Liverpool UK

**Keywords:** Bayesian modelling, gastrointestinal helminths, host-parasite interactions, parasitism, sampling regime, spatiotemporal variation

## Abstract

1. Host-parasite interactions in nature are driven by a range of factors across several ecological scales, so observed relationships are often context-dependent. Importantly, if these factors vary across space and time, practical sampling limitations can limit or bias inferences, and the relative importance of different drivers can be hard to discern.
2. We collected a replicated, longitudinal dataset of >1000 individual wood mice (*Apodemus sylvaticus*) encompassing 6 years of sampling across 5 different woodland sites to investigate how environmental, host and within-host factors determine infection intensity of a highly prevalent gastrointestinal nematode, *Heligmosomoides polygyrus.*
3. We used a Bayesian modelling approach to further quantify if and how each factor varied in space and time. Finally, we examined the extent to which a lack of spatially or temporally replication (i.e., within single years or single sites) and single (cross-sectional) versus repeated (longitudinal) sampling of individuals would affect which drivers were found to predict *H. polygyrus* infection.
4. Season, host body condition, and sex were the three most important determinants of infection intensity; however, the strength and even direction of these effects varied in time, but not in space. Additionally, longitudinal datasets, in which we can control for within-individual variation through repeat observations, provided more robust estimates of the drivers of parasite intensity compared to cross-sectional data.
5. These results highlight the importance of accounting for spatiotemporal variation in drivers of disease dynamics and the need to incorporate spatiotemporal replication when designing sampling regimes. Furthermore, they suggest that embracing rather than simply controlling for spatiotemporal variation can reveal important insight into host-parasite relationships in the wild.

## Introduction

Host-parasite dynamics in the wild are highly context-dependent, and are driven by a wide range of factors across ecological scales. These drivers range from large-scale factors such as seasonal fluctuations in environmental conditions (Nelson & Demas 1996; Dowell 2001; Altizer *et al.* 2013) or geographic variation (Davies & Pedersen 2008; Tompkins *et al.* 2011) to host-level factors such as sex (Zuk & McKean 1996), age (Plowright *et al.* 2017), or nutritional status (Sheldon & Verhulst 1996; Calder & Jackson 2000), and even include within-host effects such as interspecific interactions between co-infecting parasites or pathogens (Cox 2001; Fenton & Pedersen 2005). Importantly, these factors often vary across space and time, so their importance in driving parasitism likewise varies according to the sampling regime. Many ecological studies of infectious diseases are limited in their spatiotemporal replication and sampling breadth; hence, our ability to determine the key drivers of infection and their consistency across space and time is often limited.

Disease ecologists often seek to understand generalisable drivers of parasite dynamics, yet results are unfortunately often equivocal across studies, systems, and temporal or spatial replicates. For example, host body condition a widely used metric, often considered a proxy for fitness – is typically hypothesised to be negatively correlated with parasite infection (e.g. (Millán *et al.* 2004; Irvine *et al.* 2006; Debeffe *et al.* 2016)). However, a recent meta-analysis of >500 body condition vs. parasite infection relationships demonstrated high heterogeneity in both the strength and direction, with a high proportion (∼50%) of insignificant relationships (Sánchez *et al.* 2018). In some cases, robust sampling regimes and statistical analysis have been applied to anticipate and control for this variability. For example, a highly-replicated study of ectoparasite burden and pathogenicity in dace (*Leuciscus leuciscus*), collected over 3 years and from 8 sites, demonstrated that the effects of host and environmental factors were strong but inconsistent across years (Cardon *et al.* 2011). Even experimental approaches using parasite removal as a means of quantifying the relationships between host and parasite are vulnerable to impacts of spatiotemporal variation (Pedersen & Fenton 2015). This is highlighted by a ten-year study of red grouse (*Lagopus lagopus*), in which both host survival and reproduction (clutch size and hatching success) were greater in animals with experimentally reduced helminth burdens after drug treatment, but the magnitude of these effects varied in magnitude across years and the effects were not statistically significant in all years sampled (Hudson, Newborn & Dobson 1992). Additionally, Fenton *et al.* (2014) showed that the most reliable methods for examining drivers of parasite intensity (in this case co-infection with other parasites) involved longitudinal sampling of the same individuals, while cross-sectional approaches were less reliable. These studies make a strong case for using both well-replicated sampling and carefully structured statistical approaches to disentangle complex and context-dependent drivers of parasitism.

The regularity of finding context dependent results presents an important problem when understanding the dynamics of host-parasite interactions in the wild: specifically, what is the consistency and repeatability of the observed trends across different replicates of the same system, and how can researchers be sure that their replication is sufficient? Ecology is undergoing a reproducibility crisis, and spatiotemporal variation is a likely culprit impacting variation in results (Becker *et al.* 2020). By quantifying how the drivers of parasitism vary over space and time for the same host-parasite system, we can address three fundamental questions: (i) what are the key (reproducible) factors that drive parasite infection intensity; (ii) how much of the variation in effect sizes is biologically meaningful, rather than representing sampling biases; and (iii) how can we optimise sampling regimes to improve our ability to detect important biological variation in drivers within a system? Few ecological studies have systematically addressed these questions to date, due in part to practical sampling limitations: that is, it is difficult to sample large numbers of hosts, across a wide geographic range, and over a long period of time. Consequently, it is unclear how often wild host-parasite studies make biased inferences due to limited sampling.

European wood mice (*Apodemus sylvaticus*) are widely distributed rodents that represent an ideal system for robust spatiotemporal replication of host-parasite infection studies. They are commonly infected with an well-studied gastrointestinal nematode, *Heligmosomoides polygyrus* (Gregory 1992). Like other gastrointestinal helminths, *H. polygyrus* establishes chronic infections within their hosts, shedding eggs into the environment via faeces, where onward transmission to other host occurs after the eggs develop into infective L3 larvae (∼7 days) (Johnston *et al.* 2015). As such, *H. polygyrus* infection will be determined by the environment (Langley & Fairley 1982; Montgomery & Montgomery 1988; Gregory 1992; Brown *et al.* 1994; Abu-Madi *et al.* 2000; Eira *et al.* 2006), host factors (Gregory, Keymer & Clarke 1990; Ferrari *et al.* 2004), and within-host parasite interactions (Behnke *et al.* 2005; Knowles *et al.* 2013). Building on this knowledge, we use a temporally (6 years) and spatially (5 woodland sites) replicated wood mouse dataset to investigate the drivers of *H. polygyrus* infection intensity and their consistency across time and space. We use Bayesian modelling methods to first use the complete dataset to identify the overall drivers of infection and then to fit interactions to investigate whether these drivers varied meaningfully over space and time. Next, we use single-site or single-year subsets of the data to investigate whether the same results are found in more limited sampling regimes. Finally, we investigated the relative efficacy of cross-sectional and longitudinal sampling approaches by testing for reproducibility of the drivers of parasite infection using subsets of the data. Our approach represents a rare opportunity to quantify the drivers of parasitism and their reproducibility at multiple scales, with implications for a wide range of disease ecology studies and the design of sampling regimes.

## Methods

### Data Collection

We live-trapped several wild wood mouse populations located near Liverpool, UK regularly between May-December for six consecutive years (2009-2014). We sampled 19 trapping grids ranging in size from 2500m^2^-10,000m^2^, spread across six different woodland sites (Tables S1-2; Fig. S1). On each grid, trapping stations were placed every 10m, with two live traps (H.B. Sherman 2×2.5×6.5 in. folding traps, Tallahassee, FL, USA) at each station baited with grains and bedding material. We sampled each grid every 4 weeks from 2009-2011 and every 3 weeks between 2012-2014; with each grid trapped 3-4 nights per week. Traps were set in the late afternoon and checked the following morning. At first capture, we tagged each wood mouse with a subcutaneous microchip transponder (AVID, PIT tag), and at every capture, we recorded morphometric data (age, sex, weight, body length, reproductive condition; details below), collected a faecal sample, and examined the fur to record ectoparasite (ticks, fleas and mites) presence and intensity. Faecal samples were collected from the pre-sterilised traps, weighed and stored in 10% buffered formalin at 4°C until parasite identification.

We quantified gastrointestinal parasites (including *H. polygyrus* and coccidial protozoans *(Eimeria spp.*)) via faecal egg counts (FEC) using salt flotation to obtain the number of eggs or oocysts (for *Eimeria spp.*) per gram faeces (EPG/OPG) (Knowles *et al.* 2013). We have previously shown that EPG is highly positively correlated to adult worm burdens for *H. polygyrus* in wood mice (Sweeny *et al.* 2019). Briefly, saturated salt solution was added to formalin-preserved faecal samples so that eggs/oocysts floated to the top of the solution. The eggs were collected on a coverslip, and then we counted both eggs and oocysts at 10X magnification, and identified to species level at 40X magnification. We also identified *Eimeria spp.* to species according to unsporulated oocyst morphology (Nowell & Higgs 1989).

### Statistical Analysis

#### Defining model variables and dataset

We investigated how host-, environmental- and within-host- (eg. parasite coinfection) factors drive *H. polygyrus* infection intensity (EPG from infected animals only). In four of the six years of sampling, we conducted experiments in which randomly selected mice were anthelmintic treated or given a water control to remove/reduce gastrointestinal nematodes, such as *H. polygyrus* (see Knowles *et al.* 2013). Because anthelmintic treatment can affect *H. polygyrus* infection intensity, we restricted our analyses to only those individual mice which had not been treated. We have previously tested for, but never detected, any knock-on effects of reduced transmission or infection on untreated animals when in the presence of treated animals on the same grid. Sample sizes from all years and woodland sites can be found in Table S2.

We selected the following factors as fixed effects in all models: (i) Environmental factors: season (categorical, 3 levels: spring, summer, autumn) and (ii) Host characteristics: sex (categorical, 2 levels: male/ female); scaled mass in grams as a measure of body condition (continuous, (Peig & Green 2009)); reproductive status (categorical, 2 levels: active [males- descended or scrotal testes, females- lactating or gestating]; inactive [males- abdominal testes, females- perforate or non-perforate vagina]). Previous work from this system has demonstrated negative interactions between *Eimeria hungaryensis* and *H. polygyrus* (Knowles *et al.* 2013)), thus to investigate whether gastrointestinal (GI) parasite interactions impact infection intensity, we also included (iii) Within-host factors: the presence/absence of two most common GI parasites: the coccidian *Eimeria hungaryensis* and a Hymenolepid cestode (both categorical, 2 levels: present/absent).

#### Model Structure

Statistical analysis was carried out in R version 3.5.1, in the Bayesian linear modelling package MCMCglmm (Hadfield 2010) unless otherwise noted. MCMC methods produce a distribution of estimates for a given effect size and the proportional overlap of these estimates can be used to give a measure of significance for the difference between effects (P_MCMC_) as well as an estimate of the mean and 95% credible intervals of the difference, without the use of posthoc tests (Albery *et al.* 2018a; Palmer, Hadfield & Obbard 2018). All models were run for 130,000 iterations, with a 100-iteration thinning interval and a 30,000-iteration burnin period, for a total of 1000 stored iterations.

In order to investigate the ecological drivers of *H. polygyrus* infection intensity in wild wood mice and the impact of spatiotemporal variation in the detection of these factors, we constructed three sets of models which are described in detail below. Briefly, each model investigated the same aim – to determine what ecological factors drive *H. polygyrus* infection intensity – but contained subtle, but important differences, specifically (1) the first model set used the full data set and included both site and year as fixed effects, (2) the second model set also used the full data set but included interactions of site and year with each of the factors investigated, and lastly (3) the third model used subsets of the data set (i.e. a single site and/or year). Details for this approach are outlined below.

First (‘Model Set 1’), we used the full dataset to investigate the what factors drive *H. polygyrus* intensity across the entire 6-year, 5-woodland site study using a base model, with site and year included as fixed effects in addition to the season, host, and parasite community factors listed above. For each factor (fixed effect) we examined the proportional overlaps among the estimated posterior distributions for each site and year to determine the spatiotemporal variation of mean *H. polygyrus* intensity. Additionally, we fit an alternative model with year, site, site:year, grid, and grid:year as random effects to estimate the proportion of variance explained by each spatiotemporally relevant term.

Next (‘Model Set 2’) we added two-way interaction terms between year or site and the other fixed effect factors into the full model to investigate whether such interactions revealed spatiotemporal interactions in the environmental, host and within-host drivers of *H. polygyrus* infection intensity A visual depiction of this approach is shown in Fig. S2, where we hypothesise that unless the relationship between a factor of interest and parasite intensity is identical across space and time (Fig. S2A), estimating the variation in either the magnitude and direction of this relationship (Fig. S2B-C) will provide additional insight into parasitism in wild populations. We carried out a model addition approach using INLA (Rue, Martino & Chopin 2009) to determine the importance of the interactions without overloading the model. Starting with the base model including all fixed effects, we added interaction terms (e.g. season:year) one at a time. Each round, the interaction which lowered the DIC of the model the most (improving model fit, independent of order) was kept, and the process was repeated with the remaining interaction terms. This was repeated until the model was optimised, and could not be improved in fit with the addition of any further interaction terms (decrease DIC by >2). We ran all final model formulae in MCMCglmm. We then examined the posterior distributions of the effect estimates for interactions derived the final, optimal model, which gave an estimate for pairwise distribution overlaps for each year or site, demonstrating which years and sites differed in terms of their effect sizes for each fixed effect (e.g., the estimate of the effect of season was greater in 2010 than in 2011).

Last (‘Model Set 3’), we investigated whether different spatial and temporal sampling regimes impacted the effect size of the factors driving infection intensity, by running a series of models for either each site (N=5; with all years combined) or each year (N=6; with all sites combined). Finally, we compared how cross-sectional (single capture per individual) and longitudinal (repeat capture) sampling, defined from an individual sampling perspective (Clutton-Brock & Sheldon 2010), impacted the effect sizes of the ecological factors determining infection intensity across all three model sets. All analyses were repeated with the cross-sectional dataset which includes only the first capture for each mouse (N_i_=N_c_=783) and in a longitudinal context using all captures (N_i_= 926, N_c_=1609, max captures per individual=28, median captures per individual=4; Fig. S3). All longitudinal models included individual ID as a random effect. We report the results from the longitudinal sampling models first and then compare these models to the cross-sectional models.

## Results

### Model Set 1- Ecological drivers of parasitism with fixed spatiotemporal effects

Using our full, longitudinal (repeat captures per individual) dataset, we found that environmental (season) and within-host factors (Hymenolepid cestode coinfection) were the most important predictors of *H. polygyrus* infection intensity (Fig. 1, Table S3). Intensity was significantly lower in both summer and autumn compared to the spring (Fig. 1, P_MCMC_:Summer <0.001, P_MCMC_:Autumn <0.001) and Hymenolepid presence was associated with higher *H. polygyrus* infection intensity (Fig. 1, P_MCMC_ = 0.018). No host characteristic factors were found to significantly impact intensity of infection in this model set. In addition, we found significant spatiotemporal variation in mean *H. polygyrus* intensities across the five woodland sites and 6 years (Fig. 2). Across sites, Haddon Wood had significantly higher infection intensities compared to all other woodlands (Fig. 2A,C) and there was also significant between-year differences across several pairs of years (Fig. 2A,C). Notably, *H. polygyrus* infection intensity was higher in 2010 than in any other year, and intensities in 2012-2014 were lower than 2009-11 (Fig. 2B,D). Alternative models with spatiotemporal terms specified as random effects alongside individual ID showed that year (8.0%), individual ID (11.2%), and site (15.6%) explained the highest proportion of variance in *H. polygyrus* infection intensity; grid, grid:year, and site: year explained negligible variance (Fig. 2E).

**Fig. 1.**
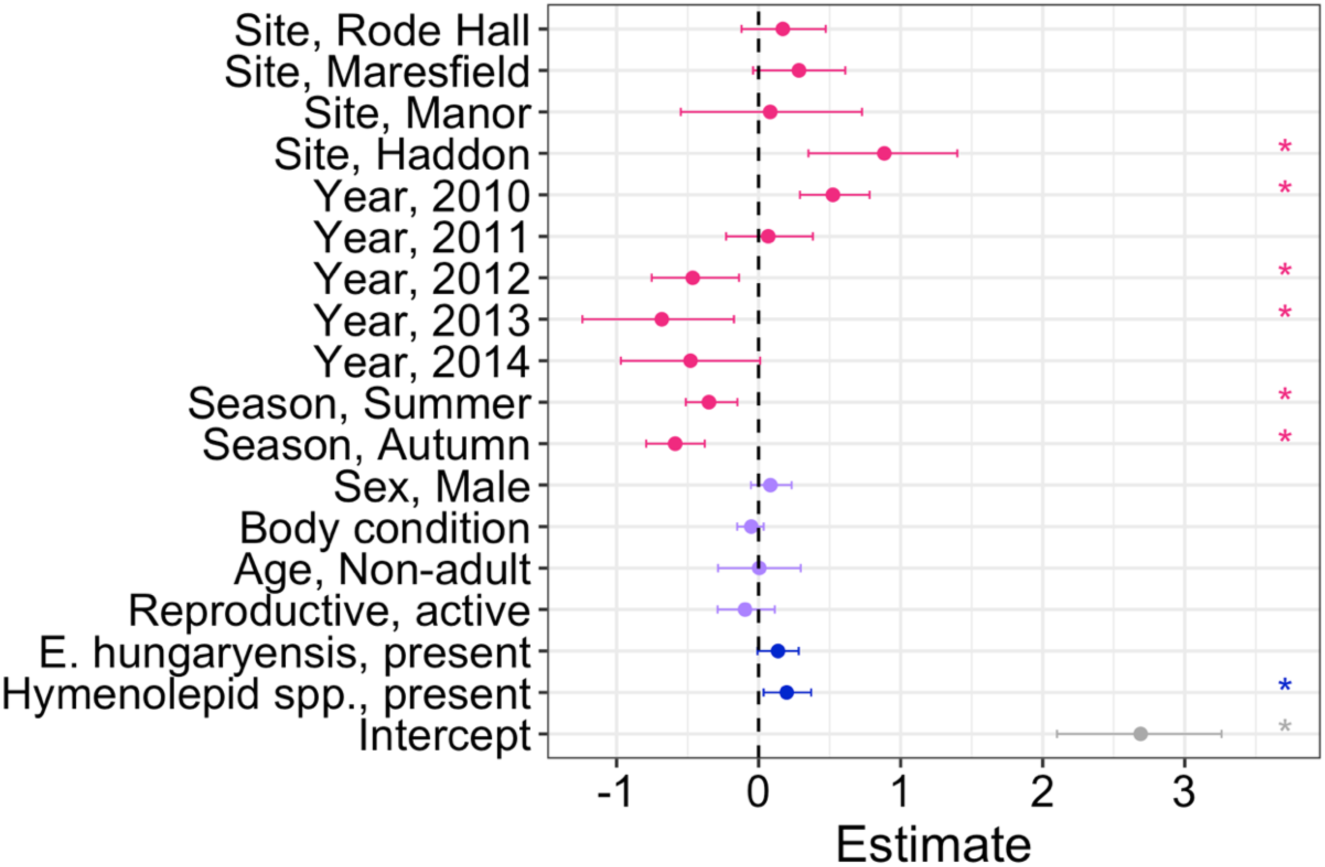
Model Set 1 output for the full dataset from all years and site collected using longitudinal sampling, where site and year were considered fixed effects. Points and ranges represent model estimates and 95% credibility estimates for each model. Colour indicates the ecological scale for each factor (pink: environment; purple: host; blue: within-host). Asterisks indicate the significance of variables with a p_MCMC_ <0.05 threshold.

**Fig. 2.**
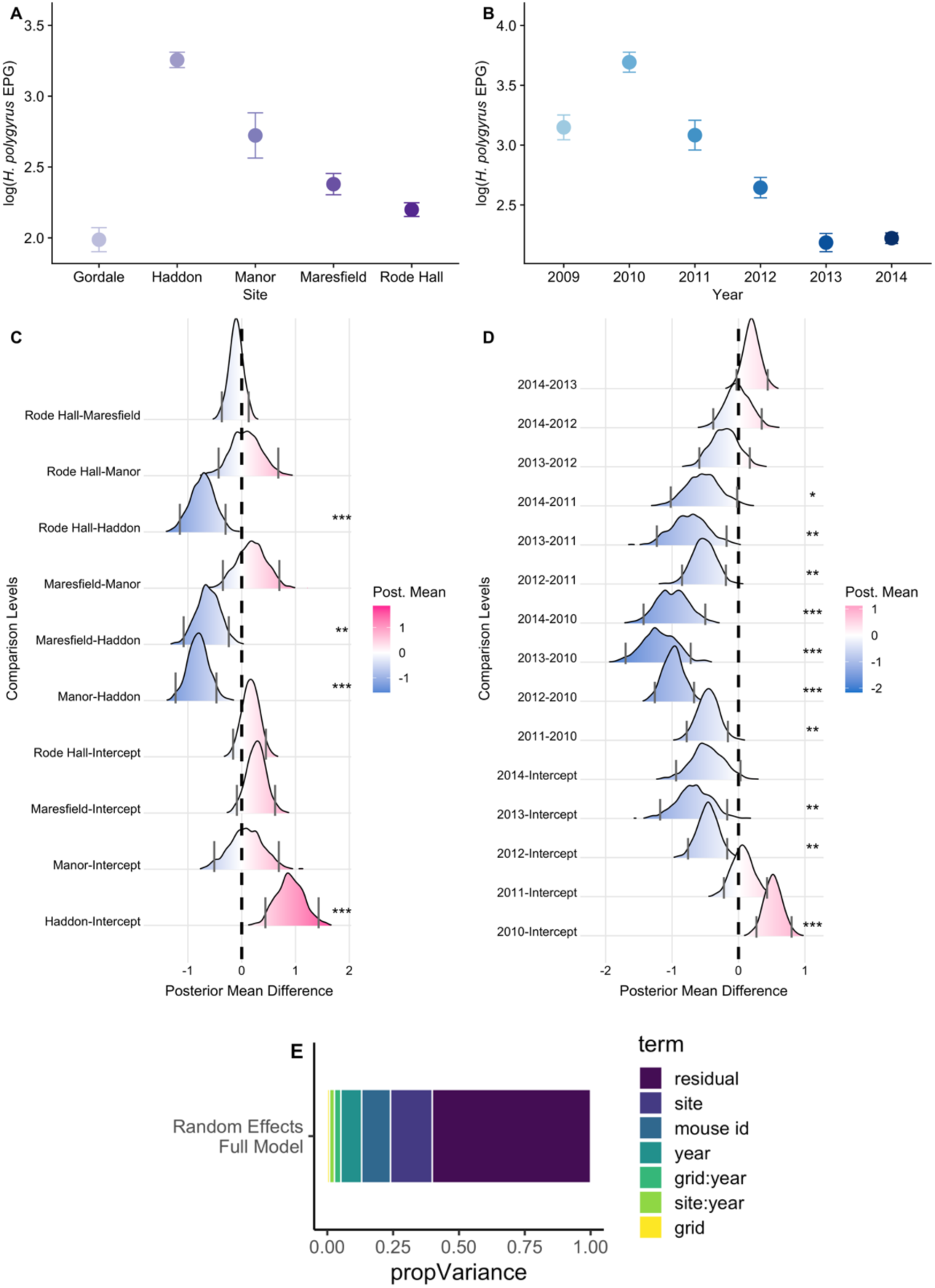
Spatiotemporal variation in mean *H. polygyrus* intensity from Model Set 1. Top row: raw data for the spatiotemporal main effects. Points represent mean intensity (± SE) for (A) site, and (B) year. Middle row: ridge plots below bar graphs represent pair-wise comparisons for the base model output for main effects of site (C) and year (D). Density ridges represent distributions drawn from the differences between the posterior means of the indicated comparison levels [a-b] for each stored iteration. Differences between effects can be interpreted by comparison of the density ridges to zero; grey lines for each ridge indicate the 95% credibility intervals for these distributions. Blue shading denotes that the mean of effect estimates for [a] is lower than that of [b] for a given interaction. Pink shading denotes that mean of effect estimates from [a] is higher than that of [b]. If credibility intervals do not cross zero, this is considered a significant difference in effects between [a-b]. Significant differences between effects are indicates by ***, ** and * for P<0.001, P<0.01 and P<0.05 respectively. ‘Intercept’ represents the baseline year of the model (2009), and Gordale for site effect levels. E. Proportion variance explained by each spatiotemporal random effect in the alternate model.

### Model Set 2: Spatiotemporal variation in ecological drivers of parasitism

Including spatiotemporal interactions with ecological factors revealed substantial temporal, but not spatial variation in the estimates for many of the factors (Fig. 3). Specifically, three factor-by-year interactions significantly improved model fit for the longitudinal models: season (ΔDIC=−58.92), body condition (ΔDIC=−14.61) and sex (ΔDIC=−5.42) (Tables S4-5; Fig. S6). Pairwise comparisons for the interactions which remained in the optimal model revealed significant changes across years for all three interactions (Fig. 3). Change in *H. polygyrus* infection intensity from Spring to both Summer and Autumn generally represented a decrease (Fig. 3B) however, comparison across years by proportional posterior overlaps still showed significant variation in both magnitude and direction of effects (Fig. 3A). The change in intensity between Spring and Summer was greatest in 2011 and 2012 (Fig. S3A-B). The decrease in intensity from Spring to Autumn for 2011-14 were greater than in 2009 and 2010, where mean intensity instead increased from Spring to Autumn (Fig. 3A-B). Interactions of year with body condition improved models by the second-largest DIC decrease (Table S5). We found a high degree of variation for the effect of body condition-by-year, where the slope between better body condition (higher scaled mass) and intensities of infection was more negative in all years 2011-2014 compared 2009, and was more negative in 2011, 2012, and 2014 compared to 2010 (Fig. 3A,D). Proportional overlap for sex-by-year interaction effects also indicated significant variation in the relationship between sex and parasite intensity. Male bias was greater in 2012-14 compared to 2010, and in 2013 compared to 2009 (Fig. 3A,C). Furthermore, we found a change in the direction of the sex-specific bias; where female wood mice had higher infection intensities than males in 2010 and 2011 (Fig. 3C).

**Fig. 3.**
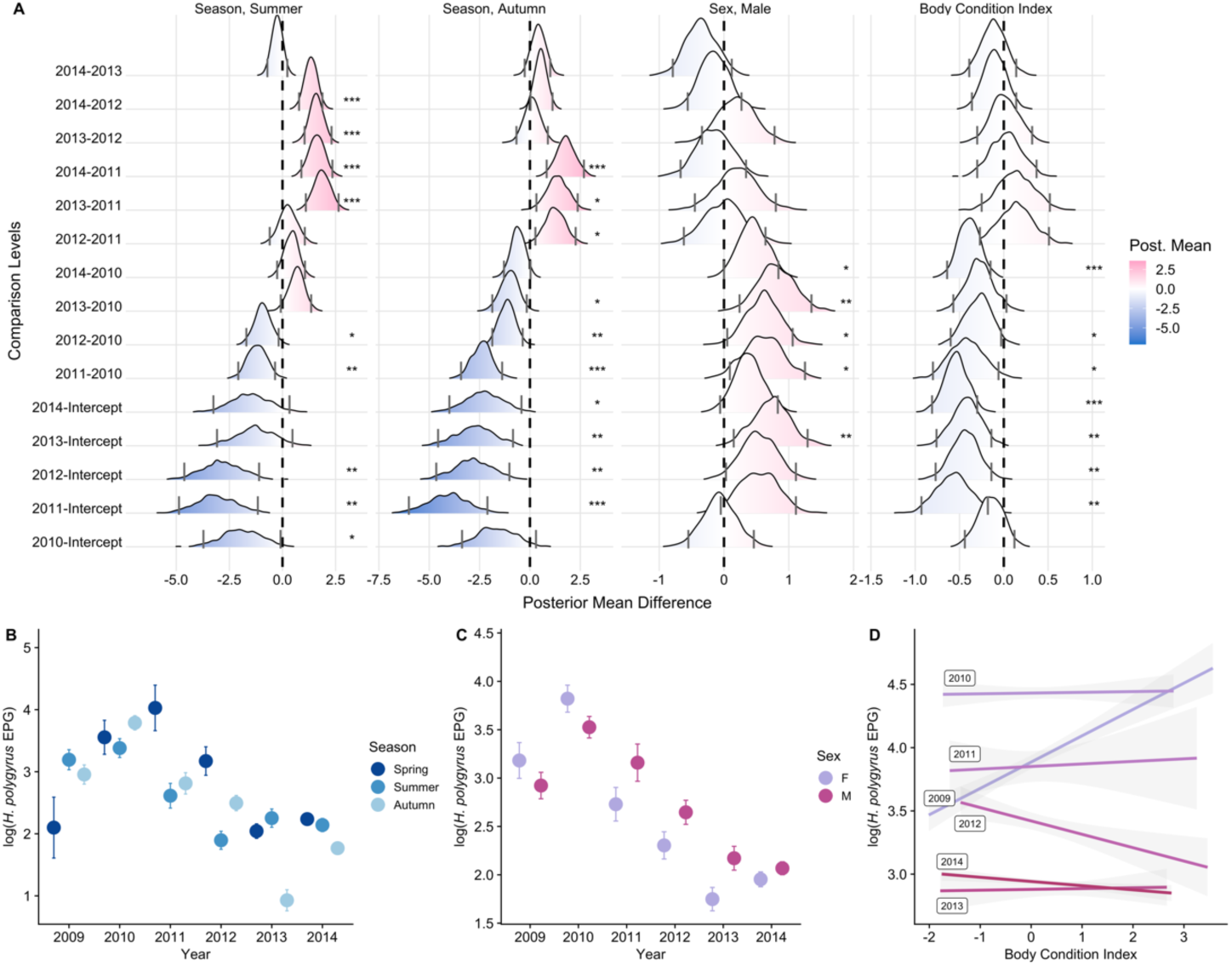
Top panel (A): Differences across estimated effects (with 95% credible intervals) for Model Set 2 including the interactions with site/year which improved full model fit for the longitudinal dataset. Density ridges represent distributions drawn from the differences between the posterior means of the indicated comparison levels [a-b] for each stored iteration. Blue shading denotes that the slope of effect from the x-axis is lower than that on the y-axis. Differences between effects can be interpreted by comparison of the density ridges to zero; grey lines for each ridge indicate the 95% credibility intervals for these distributions. Blue shading denotes that the slope of effect for [a] is lower than that of [b] for a given interaction. Pink shading denotes that slope of effect from [a] is higher than that of [b]. If credibility intervals do not cross zero, this is considered a significant difference in effect slope of [a-b]. Significant differences between effects are indicates by ***, ** and * for P<0.001, P<0.01 and P<0.05 respectively. ‘Intercept’ represents the baseline year of the model (2009) in all panels. Bottom Panel: Raw data, interactions selected for final longitudinal models (B) Season:year (C) Sex:year (D) Body condition:year. B-C represents mean +/− SE of log *H. polygyrus* intensity per category, D represents regression with SE ribbons for log *H polygyrus* intensity versus body condition index.

### Model Set 3: Ecological drivers of parasitism from limited spatial and temporal sampling regimes

We found that in models fit to data from single year or site replicates within the full longitudinal dataset, the estimates of drivers of parasite infection varied considerably (Fig. 4; Table S6-7). Across year-specific models, season was the most consistent effect in both direction and detection (Fig. 4A). Summer samples had significantly higher intensities of *H. polygyrus* infection than Spring in 2/6 years and Autumn had samples had lower intensity compared to Spring in in the majority (4/6) of year-specific models (Fig. 4A, Table S6). However, these within-year models revealed that the host characteristics were less consistent the environmental factor in their relationships with parasite intensity (Fig. 4A, Table S6). Males had higher intensity of infection compared to females in only one year, and the direction of effect was not consistent across years. Likewise, body condition had a significant association with *H. polygyrus* infection in two years, but this relationship was negative in 2014 and positive in 2009 (Fig. 4A, Table S6). The presence of both co-infecting GI parasites were associated with higher *H. polygyrus* infection intensity in just one (Hymenolepid spp.) or two (*E. hungaryensis*) years each (Fig. 4A, Table S6).

**Fig. 4.**
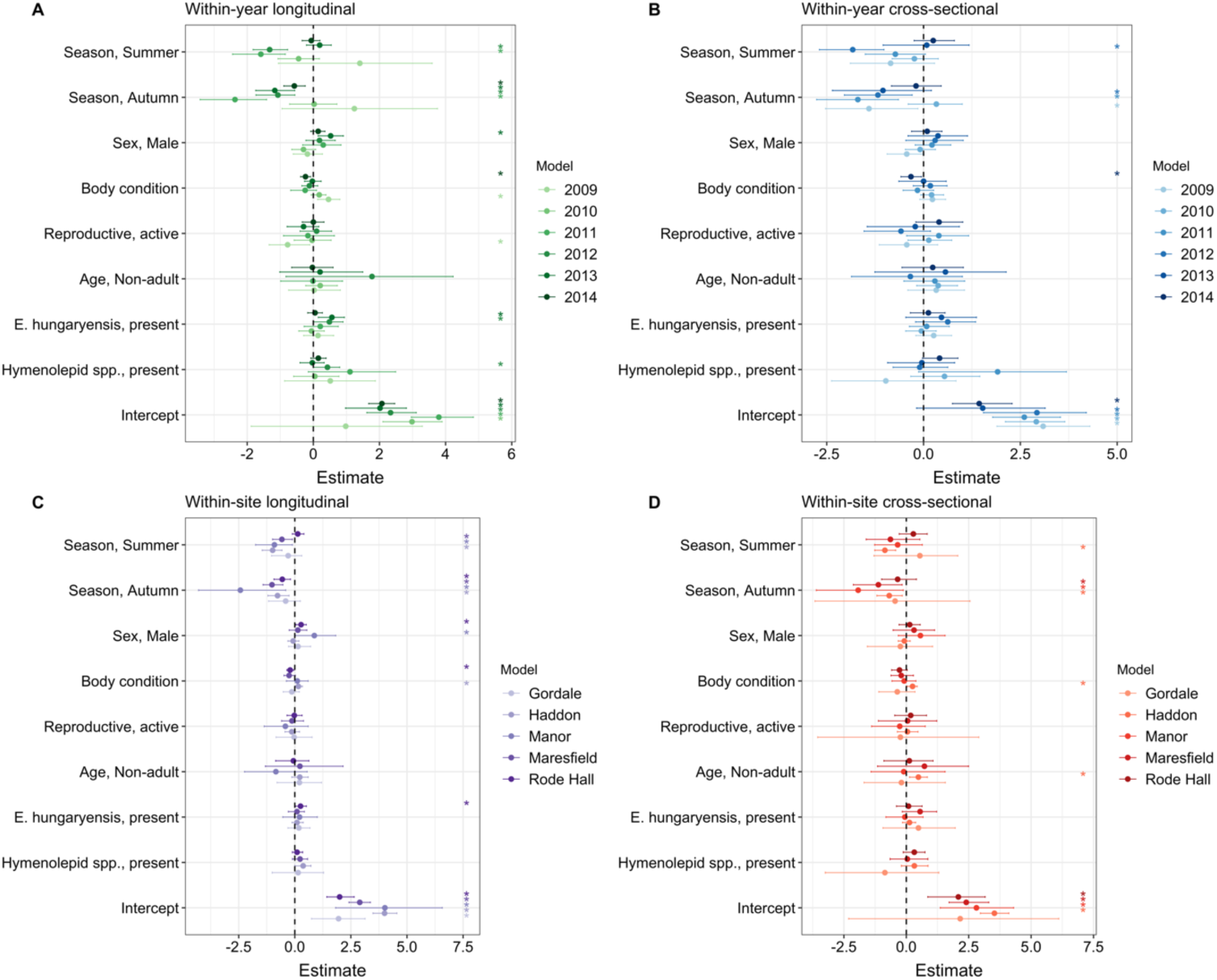
Effect size estimates from Model Set 3, investigating the effect environment, host, and parasite community-level predictors of *H. polygyrus* intensity within longitudinal (A,C) and cross-sectional (B,D) sampling resolutions of years (A,B) and sites (C,D). Points and ranges represent model estimates and 95% credibility estimates for each model. Asterisks indicate significance at a threshold of P<0.05. Intercepts for categorical covariates are as follows: Season, Spring; Sex, Female; Reproductive status, Inactive; Age, Adult; *E. hungaryensis* & Hymenolepid spp., absent.

In the five site-specific models, we found again that the seasonal effects were the most consistent and frequently detected (Fig. 4C, Table S7). Specifically, Summer samples had lower infection intensities than Spring for 3/5 sites and Autumn in 4/5 sites. Body condition had a significant relationship with parasite intensity but in 2/5 sites, and the direction of this effect varied. Similarly, we found a significant male bias in only 2/5 site. Lastly, we found positive associations between the presence of *E. hungaryensis* and *H. polygyrus* intensity for one woodland site only (Fig. 4, Table S7).

### Comparison of cross-sectional versus longitudinal sampling

All results described above used the longitudinal sampling dataset, which included multiple sampling points per individual. When we ran the same analyses using just cross-sectional data (a subset of the longitudinal dataset with only a single sample point (first capture) for each mouse) we found qualitatively similar results but less confidence in estimated effects. For the cross-sectional models of the full dataset without interactions (Model Set 1), all effects were estimated with wider 95% credibility intervals than in longitudinal models (Fig. S4). For Model Set 2, cross-sectional models and longitudinal models differed in regard to both the year and site interactions which improved model fit (Fig. S6; Table S5). Although cross-sectional models were improved with the same interactions retained in optimal longitudinal models, their posterior overlaps for interaction detected less variation than those from the corresponding longitudinal optimal model (Figs 3;S6-7). Furthermore, cross-sectional models were also improved by three additional interactions compared to the corresponding longitudinal model (Table S5): sex-by-site, reproductive status-by-year, and Hymenolepid presence-by-year. However, there was very little variation across these interaction levels (Fig. S7); only the effect of Hymenolepid presence varied significantly across interaction levels, where Hymenolepid presence in 2011 had a stronger positive association with *H. polygyrus* intensity than in all other years.

Among models fit to data subsampled within each year or site (Model Set 3), cross-sectional models largely agreed with the direction of effect compared to longitudinal models, but detected fewer significant effects for all ecological scales considered (environment, host, and parasite community) (Fig. 4). As with longitudinal sampling models, effects of season were largely consistent in cross-sectional models, but effects of host factors and co-infecting parasite presence were detected less frequently, and variations detected by interactions in optimal interaction models for the full dataset were estimated with lower confidence (wider 95% credibility intervals; Fig. 4).

## Discussion

Using an extensively replicated dataset, we demonstrate that drivers of parasitism can vary across space and time in a wild mammal population, and that the observed relationships depend intimately on the sampling effort used to detect them. Specifically, we give evidence for consistent overarching effects of seasonality, context-dependent effects of sex and body condition, and varying detectability of within-host coinfection associations (Table 1). These results suggest that ecological drivers of parasitism are not likely to be static, and highlight the importance of examining spatiotemporal context when estimating drivers of parasitism, encouraging caution when generalising from analyses that are limited in their spatial or temporal replication.

**Table 1.** Comparison of the ecological factors driving *H. polygyrus* infection intensity across the 3 models sets.

| Fixed Effect | Cross-sectional |  |  | Longitudinal |  |  | Implication |
| --- | --- | --- | --- | --- | --- | --- | --- |
|  | Model Set 1<br>Base Model | Model Set 2<br>With<br>Interactions | Model Set 3<br>Submodels | Model Set 1<br>Base Model | Model Set 2<br>With<br>Interactions | Model Set 3<br>Submodels |  |
| Season, Summer | Y | Y* | Y | Y | Y* | Y | Seasonality impacts parasitism across all spatiotemporal scales, but the slopes of these effects vary between years |
| Season, Autumn | Y | Y* | Y | Y | Y* | Y |  |
| Sex, Male | N | Y* | N | N | Y* | Y | No **overall** effect of host characteristics driving infection intensity, but significant variation in the slopes and direction of effects in interaction models & subsampled models, which indicates context-dependency in these relationships |
| Body condition | N | Y* | Y | N | Y* | Y |  |
| Reproductive, active | N | Y* | N | N | N | N |  |
| *E. hungaryensis* present | N | N | N | Y | N | Y | Overall, a positive association between co-infecting GI parasites and *H. polygyrus* with little significant variation year-to-year but significant variation in detection within single years and site replicates |
| Hymenolepid present | N | Y* | N | Y | Y | Y |  |

Blue shading ('Y') indicates a significant relationship between the given fixed effect and *H. polygyrus* intensity, while a 'N' suggests no relationship. For Model Set 2, asterisks indicate a significant interaction between a given factor and year or site. Implications give a broad-level synthesis of discussion presented in the main text.

Despite large sample sizes, our full dataset (Model Set 1) revealed surprisingly weak overarching evidence for host effects on parasitism, with much stronger support for environmental effects as key drivers. Aside from weak positive correlations between both coinfecting GI parasites (*Eimeria hungaryensis* and Hymenolepid) presence, and *H. polygyrus* infection intensity, no host-level characteristics had significant effects across all years and sites. This somewhat contradicts results from several smaller scale studies of *H. polygyrus* in *Apodemus spp.* populations, many of which document significant host-level drivers such as male-bias in both transmission (Ferrari *et al.* 2004) and burden (Langley & Fairley 1982; Gregory *et al.* 1990). Similarly, effects of age (Behnke *et al.* 1999; Clerc *et al.* 2019), reproduction (Albery *et al.* 2018b), body condition (Warburton, Pearl & Vonhof 2016), and interactions between co-infecting parasites (Behnke *et al.* 2005; Clerc *et al.* 2018) are well-documented in mammal-helminth systems. Instead, the strongest and most consistent drivers were the environmental factors of year, site, and season, agreeing with extensive previous work in multiple *A. sylvaticus* populations (Montgomery & Montgomery 1990; Gregory 1992; Gregory, Montgomery & Montgomery 1992; Abu-Madi *et al.* 1998; Behnke *et al.* 1999; Abu-Madi *et al.* 2000; Eira *et al.* 2006; Bordes *et al.* 2012). Taken together, these findings imply that environmental drivers across space and time dwarf the effects of individual host-level factors, and may even prevent their detection so that within-site studies may be better-suited to host factor-level analyses. To investigate this possibility, we asked: did additional drivers of *H. polygyrus* infection vary significantly in their effects across sites and years, so that aggregating data across the whole study conceal this variation?

Our interaction analysis (Model Set 2) revealed considerable variation in season, sex, and body condition effects across different time and/or space. Across many helminth species, abundance is often highest in the spring and declines throughout the winter (Stromberg 1997), and our results from our initial model (Model Set 1), support these seasonal patterns. Our interaction models, however, show that the magnitude and direction of this effect changes significantly across years, and in some years intensities in Spring are in fact lower than Summer or Autumn. In the case of host sex and body condition, our interaction models revealed significant relationships between both factors and *H. polygyrus* intensity that were masked when data was aggregated across years. This aligns with fundamental observations of host factors as important drivers of parasitism across a range of systems (Zuk & McKean 1996; Sánchez *et al.* 2018), but indicates that temporally varying factors can alter their relationships with helminth infection. There are several possible reasons for such temporal variation. For example, seasonality in helminth dynamics can be linked to changing hostage structures (Keymer & Dobson 1987). Wood mouse population dynamics can vary substantially from year to year, which may have altered seasonal fluctuations of helminth infection. Likewise, cross-species meta-analyses of both sex bias (Moore & Wilson 2002) and body condition effects (Sánchez *et al.* 2018) have shown that both effects can vary by host system and/or the method of sampling. Our results corroborate this observation within a single host species and imply that the detection of these relationships is dependent on sampling efforts. Next, we asked: were single-site or single-year models were capable of detecting these effects,

Subsetting our analysis into single-site and single-year datasets (Model Set 3) revealed substantially weaker models and some likely spurious results. By combining the findings from these models with the more powerful across-site models incorporating interaction effects (Model Set 2), we were able to show that limited sampling replication may produce misleading results (Table 1). For example, results from 2009 alone indicate no seasonal effects on *H. polygyrus* intensity, lower intensities of infection in reproductively active individuals, and higher intensities in individuals with higher body condition scores—results which are not supported by results from any other single year or site subset or by models using the full dataset. These results suggest that caution should be taken when inferring biological relationships from limited spatial or temporal sampling scales. To further investigate the impact of sampling regime datasets on the drivers of parasite infection and investigate options for maximizing interpretation of results even when spatiotemporal replication is limited, we finally compared results models from both the full dataset and years and site subsets with their cross-sectional correlates in which only one observation per individual to ask how sampling design influences model estimates.

There was universal support for longitudinal models outperforming cross-sectional in all models included in this study, both in terms of model certainty and the detection of biologically meaningful results. In the interaction models (Model Set 2), cross-sectional model selection retained the same terms as in the longitudinal models, but also retained additional interaction effects (eg. Reproductive status & Hymenolepid presence). However, these models detected fewer significant differences across interaction levels for those interactions retained in both longitudinal and cross-sectional models, and very little signification variation for the interactions retained only in cross-sectional models (Fig. S7). This lack in detection of significant differences across levels for the additional interactions maintained in the cross-sectional models suggest that these were likely spurious, originating from reduced power rather than real biological variation. For models within single years or sites (Model Set 3), longitudinal models similarly revealed more precise estimation (narrower confidence intervals) of effects for key drivers of *H. polygyrus* intensity that were also identified in Model Sets 1 and 2 using the full dataset. These differences between cross-sectional and longitudinal models are notable given that the median number of observations per individual mouse in our dataset is low (4) and very few animals approach the maximum number of recorded sampling points (28; Fig S3). This suggests that the benefit of longitudinal sampling for estimating drivers of parasitism is conferred even with very few additional data points per individual. These findings generally agree with studies revealing that longitudinal sampling is more robust for detecting parasite interactions (Fenton *et al.* 2014), and highlight the broad benefits of longitudinal sampling for multiple areas of interest within wildlife ecological studies particularly where replicates across space and time are not possible. Encouragingly, however, cross-sectional effect estimates did largely follow the trends of longitudinal (Fig. S4), indicating that cross-sectional sampling can provide reliable insight into data trends as long as observed effects are not overinterpreted.

Although our sampling regime was not suited to identifying specific environmental drivers such as climatic or resource availability, most variation occurred at the between-year rather than the between-site level, implying that within this system temporal (and not spatial) variation in both environmental and host factors driving *H. polygyrus* intensity is more important. However, year and site combinations in this dataset were not perfectly independent, and so it is difficult to fully disentangle the impact of space form time (Table 1). It is therefore possible that some apparent temporal variation is attributable to spatial site changes, and *vice versa*. In addition, the woodlands included in this study were initially chosen because they were similar deciduous woodlands, and it is therefore possible that the habitats were not diverse enough to detect biological variation in drivers of parasitism due to spatial environmental context. Given previous work showing variable effects of seasonality on helminth infections of *A. sylvaticus* across highly different habitats (i.e. sand dunes versus lake margins) (Eira *et al.* 2006), further application of this approach to include datasets across varied habitats and addressing more fine-scale spatial variation would therefore be useful. This study also focused exclusively on those factors that influence infection intensity and their spatiotemporal variation. Although helminth egg count was examined here and is a common proxy for infection burden (Margolis *et al.* 1982), prevalence (proportion of individuals infected in a population) is also a key metric of disease for many study systems and it is worth nothing that factors that increase exposure and infection probability may therefore differ from those that dictate infection intensity, and this should be kept in mind for interpretation of these results.

By carrying out substantial spatiotemporal sampling replication, we have provided evidence for important temporal changes in seasonality, sex, and body condition effects on helminth infection dynamics in a wild wood mouse population. As well as confirming significant spatiotemporal variation in parasitism itself, our findings suggest that the drivers of infection intensity exhibit variation over space and time. This suggests that varied support for hypotheses regarding factors influencing parasitism may represent important biological variation rather than simply variation in detection or limited statistical power across different populations and replicates. Given practical limitations of many ecological studies, it is typically of interest to determine whether sampling resolution in a wild system alters estimates of effects of interest. Our results suggest that longitudinal sampling within limited replicates provides reliable estimates of effects, but highlight that caution should be applied in extrapolating these results beyond the context of the study. Overall, we hope that more studies will investigate and control for spatiotemporal variation in effects influencing important aspects of host-parasite dynamics.

## Data accessibility

Data will be made available from the Dryad Digital Repository upon acceptance.

## Acknowledgements

This work was supported by PhD studentships from the Darwin Trust of Edinburgh to ARS and SV. A.F. and A.B.P. are supported by grants from the National Environment Research Council (NERC; NE/ I024038/1 and NE/I026367/1) and A.B.P. is funded by a Chancellors Fellowship and a Wellcome Trust Strategic Grant for the Centre for Immunity Infection and Evolution (Grant 095831). We thank the many tireless field assistants and volunteers involved in field and laboratory data collection. Lastly, we thank the EGLIDE (Edinburgh, Glasgow, and Liverpool Infectious Disease Ecology) group for their feedback and help with the analysis and interpretation of our results.

## Author Contributions

Data for this study was provided by long-term study led & designed by ABP & AF. ARS conceived of the analysis with ABP & AF. ARS, GFA, & SV designed statistical approach. ARS analysed the data. ARS led the writing of the manuscript. All authors contributed critically to analysis discussion and manuscript drafts.

## Supplementary Information

**Figure S1.**
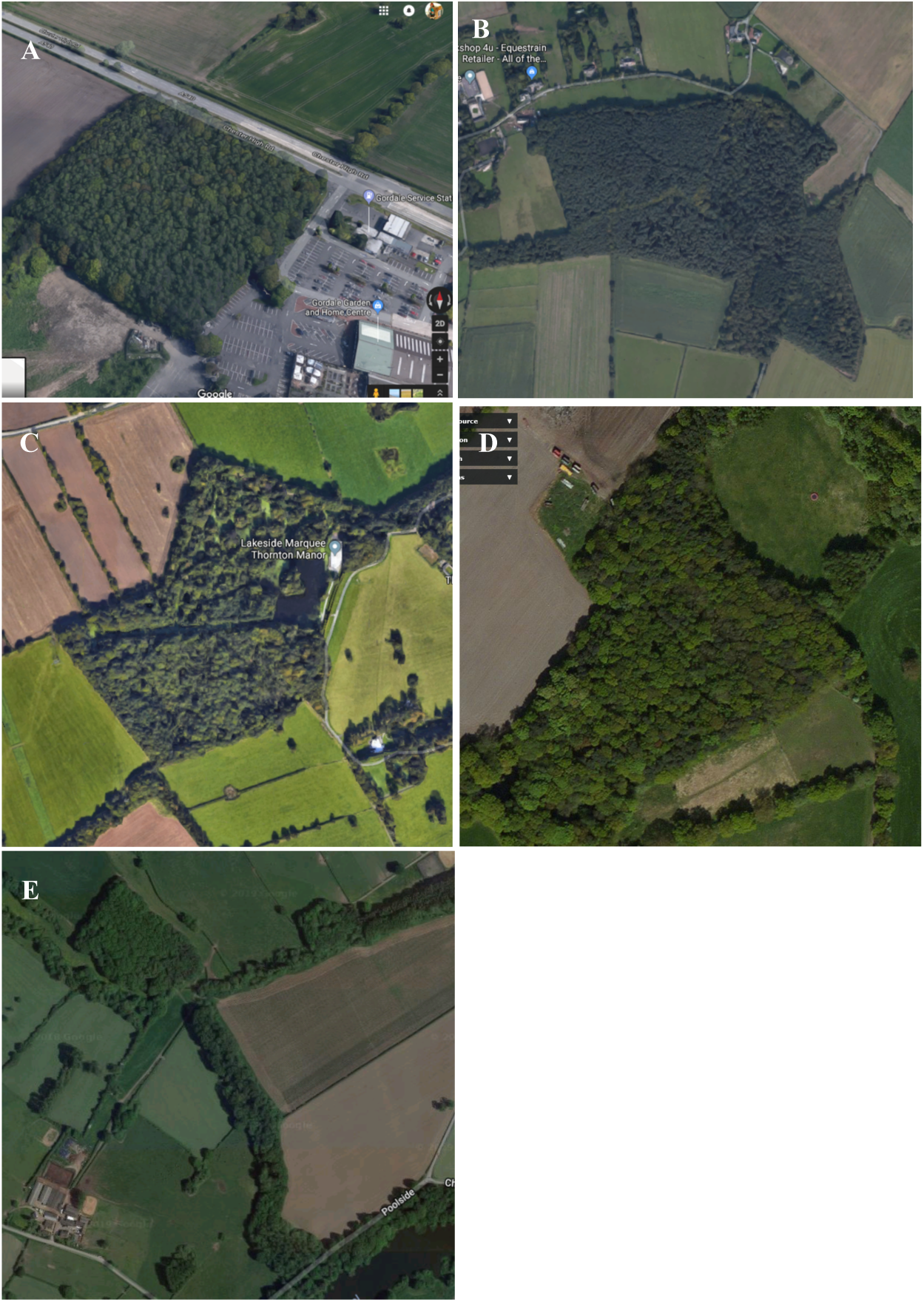
Sites included in dataset. A. Gordale B. Haddon Wood C. Manor Wood D. Maresfield Farm E. Rode Hall

**Table S1.**
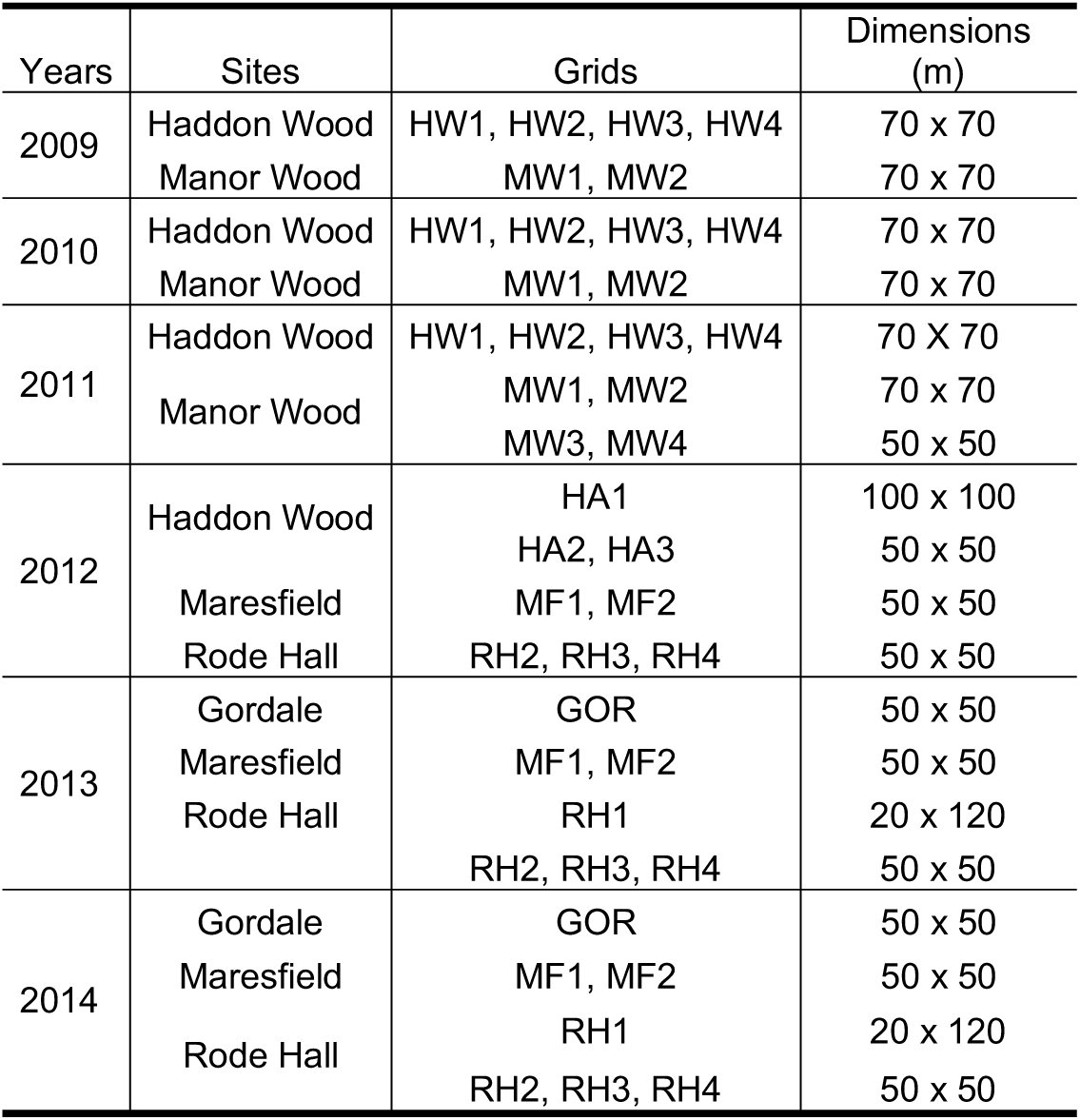
Dimensions of grids included in analysis.

**Table S2.** Sample size and prevalence of *H. polygyrus* across all site-year combinations.

|  | Site |  |  |  |  |  |  |  |  |  | Total<br>(N, Inf) |
| --- | --- | --- | --- | --- | --- | --- | --- | --- | --- | --- | --- |
|  | Gordale |  | Haddon Wood |  | Manor Wood |  | Maresfield |  | Rode Hall |  |  |
|  | N | Prevalence (%) | N | Prevalence (%) | N | Prevalence (%) | N | Prevalence (%) | N | Prevalence (%) |  |
| 2009 |  |  | 317 | 56 | 82 | 9 |  |  |  |  | 399 (185) |
| 2010 |  |  | 327 | 64 | 104 | 14 |  |  |  |  | 431 (222) |
| 2011 |  |  | 228 | 39 | 168 | 21 |  |  |  |  | 396 (124) |
| 2012 |  |  | 581 | 30 |  |  | 239 | 15 | 149 | 18 | 969 (241) |
| 2013 | 115 | 9 |  |  |  |  | 214 | 16 | 852 | 19 | 1181 (206) |
| 2014 | 404 | 27 |  |  |  |  | 794 | 21 | 1174 | 31 | 2372 (631) |
| Total<br>(N, Inf) | 519 (117) |  | 1453 (652) |  | 354 (56) |  | 1247 (236) |  | 2175 (548) |  | 5748 (1609) |

Blank cells represent instances where sites were not sampled in a given year. Ns represent total observation for a given category; data used for all models included infected individuals only (percentage of total indicated by prevalence). Total values in parentheses indicate totals for infected individuals in final analyses.

**Figure S2.**
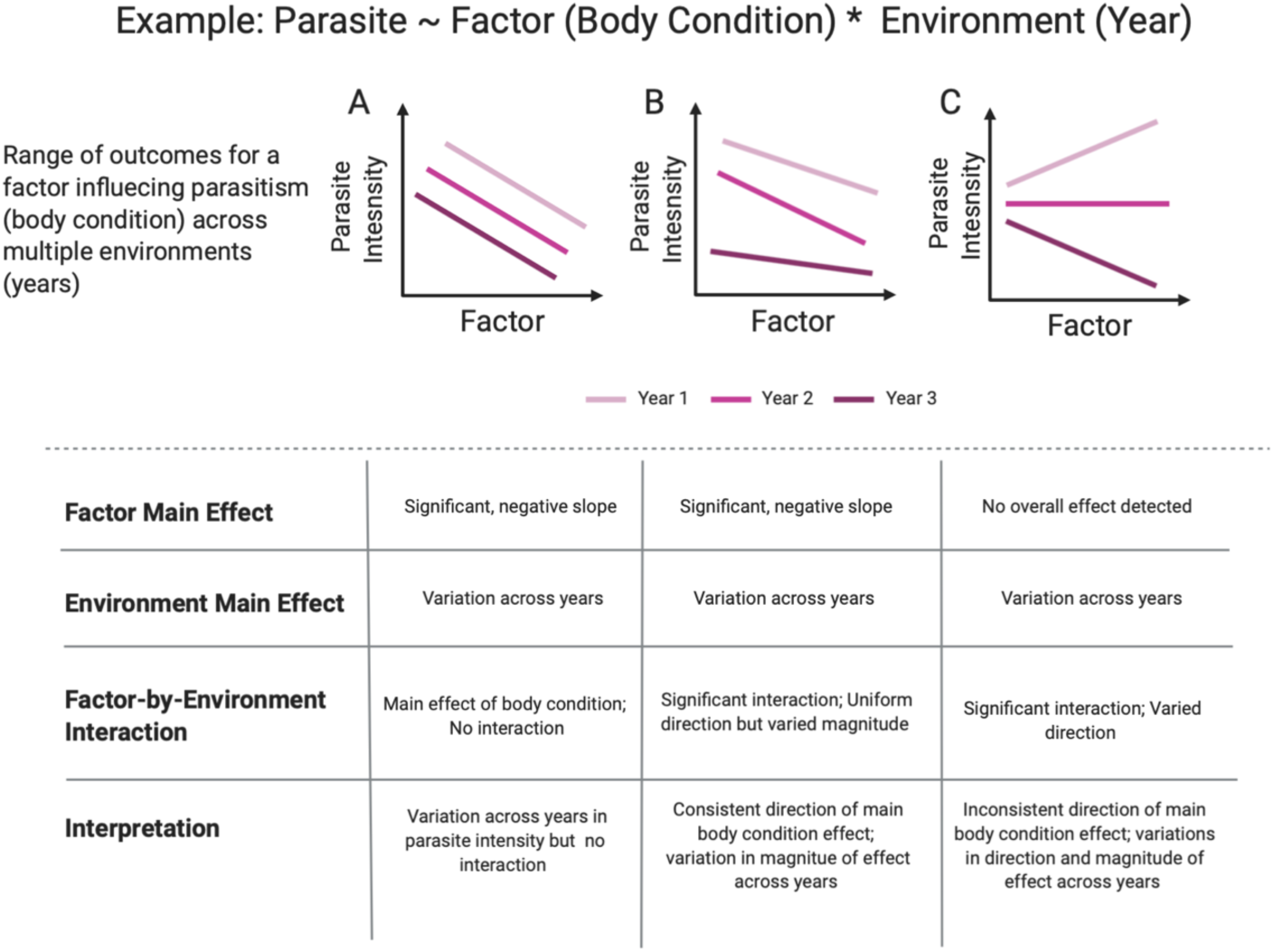
Statistical approaches for the analysis of the drivers of infection intensity across multiple environments (eg. study year or site). An illustrative example of the relationship between body condition and parasite intensity across three years is presented, with implications for each scenario.

**Figure S3.**
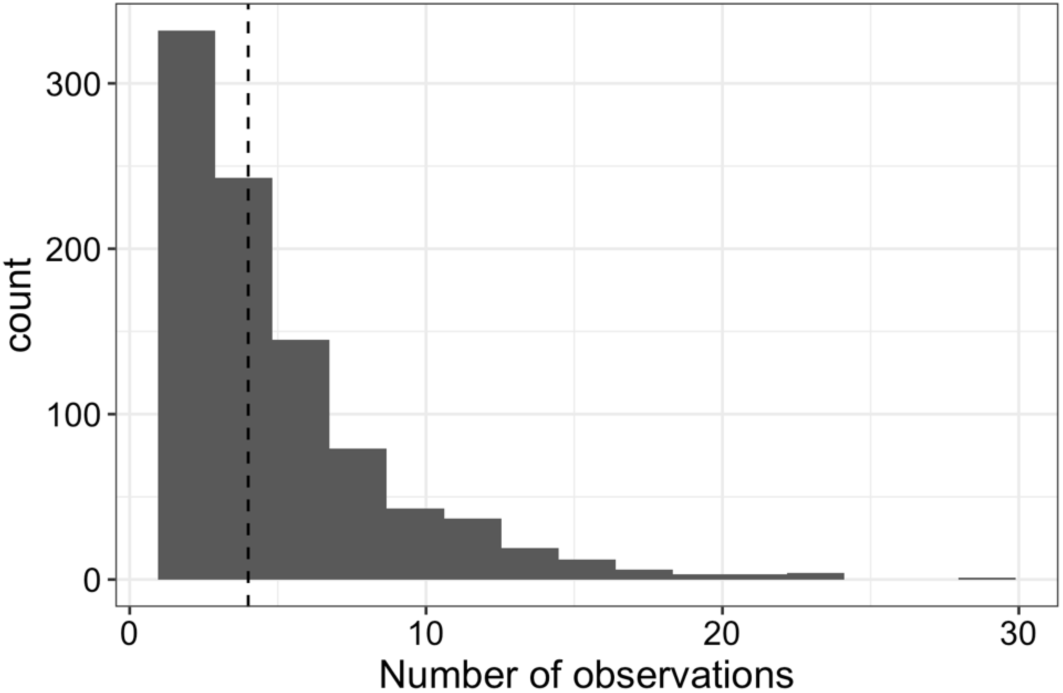
Histogram of observations per individual included in longitudinal models. N_i_=783; N_c_=1609. Dashed line represents the median number of observations.

**Table S3.** Full Model Output, without interactions.

| variable | Cross-sectional |  | Longitudinal |  |
| --- | --- | --- | --- | --- |
|  | Estimate | P<sub>MCMC</sub> | Estimate | P<sub>MCMC</sub> |
| Site, Rode Hall | 0.13 (-0.43 - 0.67) | 0.646 | 0.17 (-0.16 - 0.45) | 0.294 |
| Site, Maresfield | 0.31 (-0.28 - 0.93) | 0.342 | 0.27 (-0.09 - 0.62) | 0.118 |
| Site, Manor | 0.05 (-0.92 - 1.1) | 0.924 | 0.09 (-0.51 - 0.69) | 0.778 |
| Site, Haddon | 0.83 (-0.04 - 1.78) | 0.09 | 0.9 (0.44 - 1.43) | 0.001 |
| Year, 2010 | 0.63 (0.35 - 0.93) | 0.001 | 0.52 (0.27 - 0.8) | 0.001 |
| Year, 2011 | 0.23 (-0.12 - 0.56) | 0.188 | 0.06 (-0.22 - 0.43) | 0.698 |
| Year, 2012 | -0.26 (-0.69 - 0.13) | 0.22 | -0.46 (-0.76 - -0.17) | 0.004 |
| Year, 2013 | -0.67 (-1.65 - 0.09) | 0.122 | -0.67 (-1.18 - -0.17) | 0.01 |
| Year, 2014 | -0.37 (-1.23 - 0.41) | 0.362 | -0.47 (-0.94 - 0.03) | 0.058 |
| Season, Summer | -0.32 (-0.57 - -0.02) | 0.02 | -0.35 (-0.54 - -0.19) | 0.001 |
| Season, Autumn | -0.49 (-0.83 - -0.15) | 0.006 | -0.6 (-0.81 - -0.35) | 0.001 |
| Sex, Male | -0.04 (-0.25 - 0.13) | 0.666 | 0.09 (-0.05 - 0.23) | 0.23 |
| Body condition | 0.02 (-0.11 - 0.18) | 0.788 | -0.05 (-0.15 - 0.04) | 0.288 |
| Age, Non-adult | 0.28 (-0.03 - 0.56) | 0.056 | 0 (-0.29 - 0.28) | 0.982 |
| Reproductive, active | 0.11 (-0.18 - 0.41) | 0.472 | -0.1 (-0.28 - 0.09) | 0.306 |
| *E. hungaryensis*, present | 0.11 (-0.1 - 0.32) | 0.282 | 0.14 (0.01 - 0.29) | 0.058 |
| Hymenolepid, present | 0.2 (-0.13 - 0.51) | 0.224 | 0.2 (0.04 - 0.37) | 0.018 |
| Intercept | 2.36 (1.39 - 3.37) | 0.001 | 2.69 (2.1 - 3.26) | 0.001 |

Shading represents significant effects at threshold $P_{\text{MCMC}} < 0.05$ .

**Figure S4.**
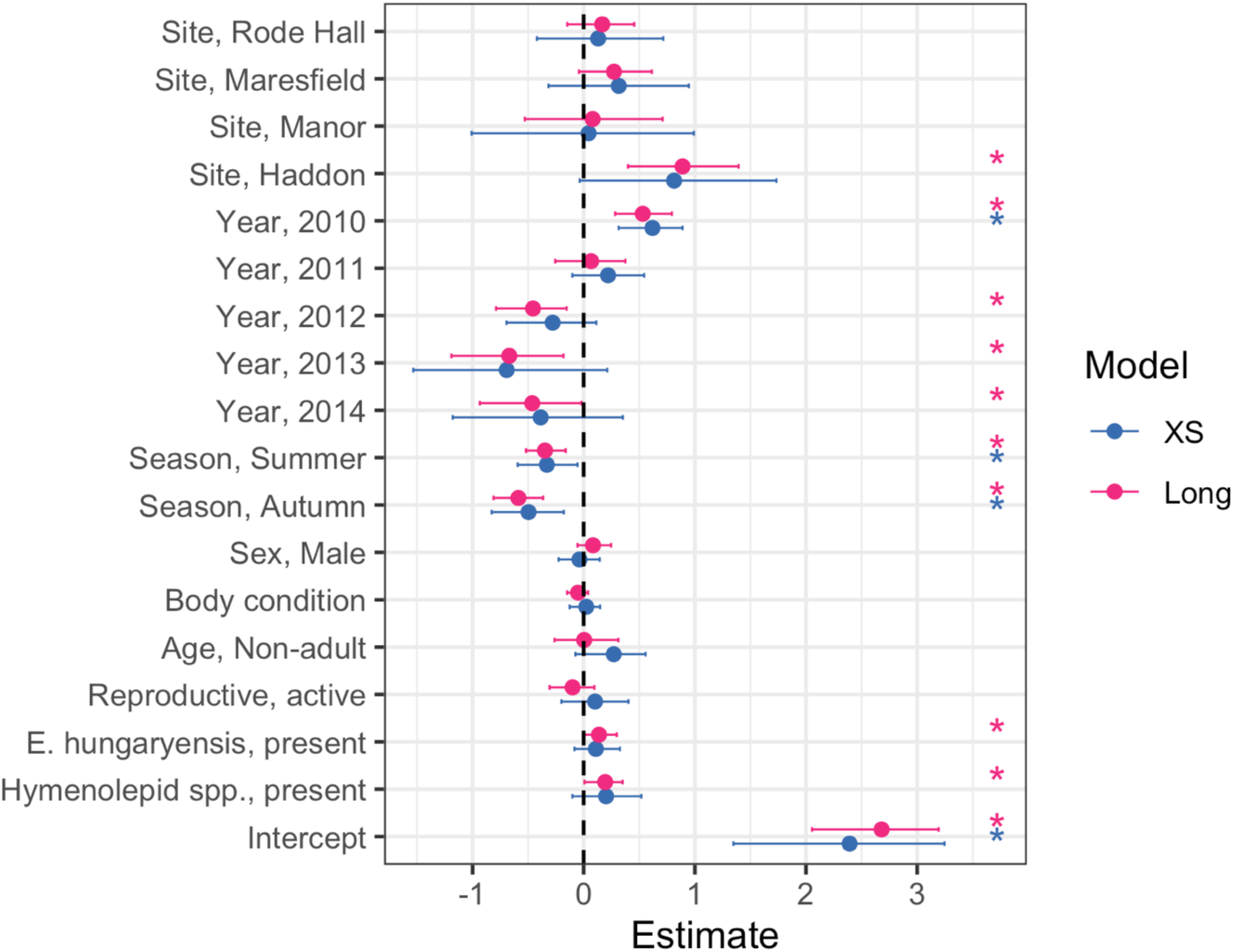
Full model output for cross-sectional and longitudinal models on data from all years and sites. Points and ranges represent model estimates and 95% credibility estimates for each model. Asterisks indicate the significance of variables with a p_MCMC_ <0.05 threshold.

**Figure S5.**
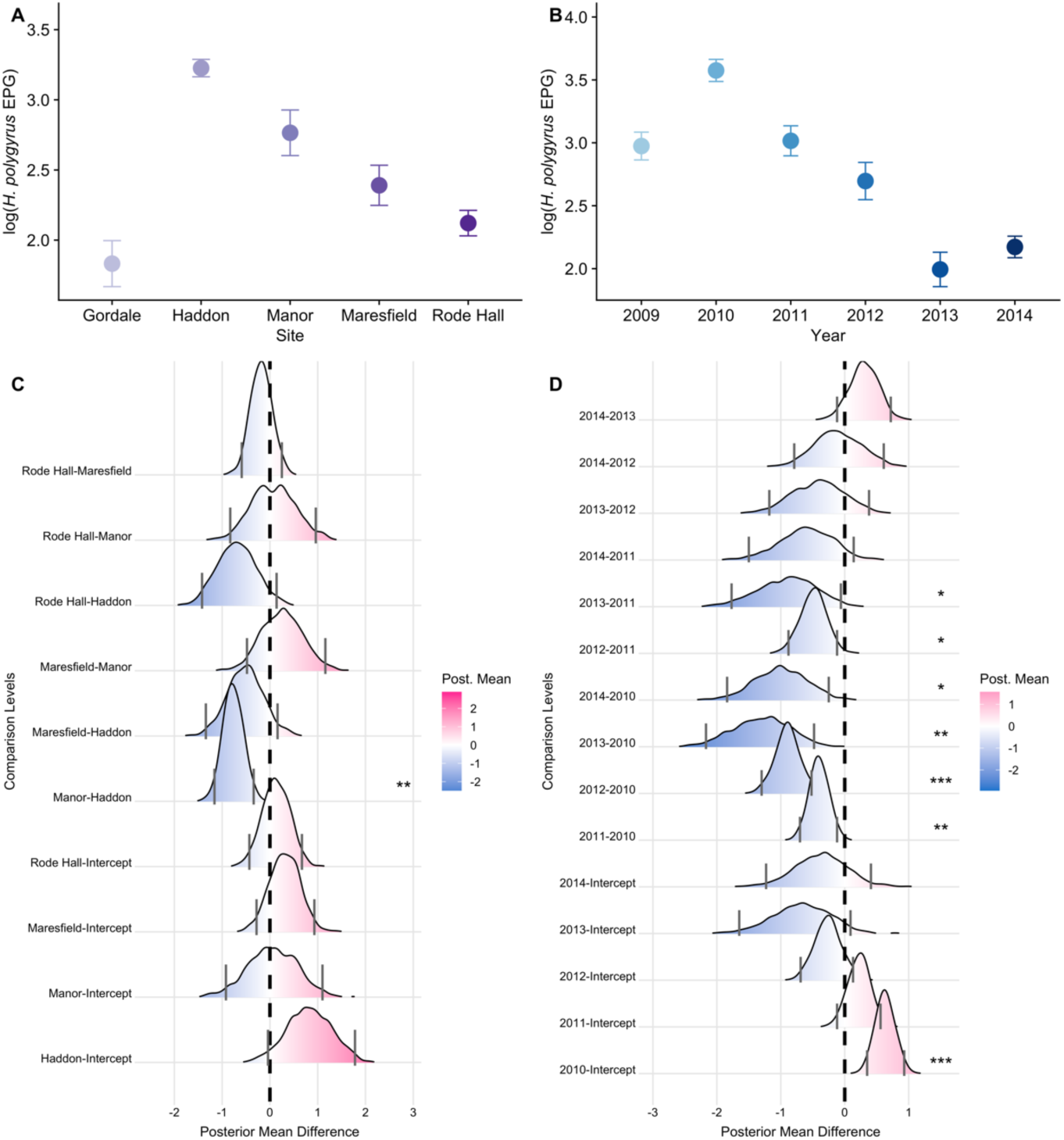
Spatiotemporal variation in mean *H. polygyrus* intensity across cross-sectional dataset. Raw data for significant spatiotemporal main effects from cross-sectional base model: Points represent mean intensity (± SE) for A. site and B. year. Ridge plots below bar graphs (C-D) represent the pair-wise comparisons for base model output for fixed effect factors. Density ridges represent distributions drawn from the differences between the posterior means of the indicated comparison levels [a-b] for each stored iteration. Blue shading denotes that the mean of effect estimates from the x-axis is lower than that on the y-axis. Differences between effects can be interpreted by comparison of the density ridges to zero; grey lines for each ridge indicate the 95% credibility intervals for these distributions. Blue shading denotes that the mean of effect estimates for [a] is lower than that of [b] for a given interaction. Pink shading denotes that mean of effect estimates from [a] is higher than that of [b]. If credibility intervals do not cross zero, this is considered a significant difference in effects between [a-b]. Significant differences between effects are indicates by ***, ** and * for P <0.001, P <0.01 and P <0.05 respectively. ‘Intercept’ represents the baseline year of the model (2009) in all panels. ‘Intercept’ represents Gordale for site effect levels and ‘2009’ for year.

**Figure S6.**
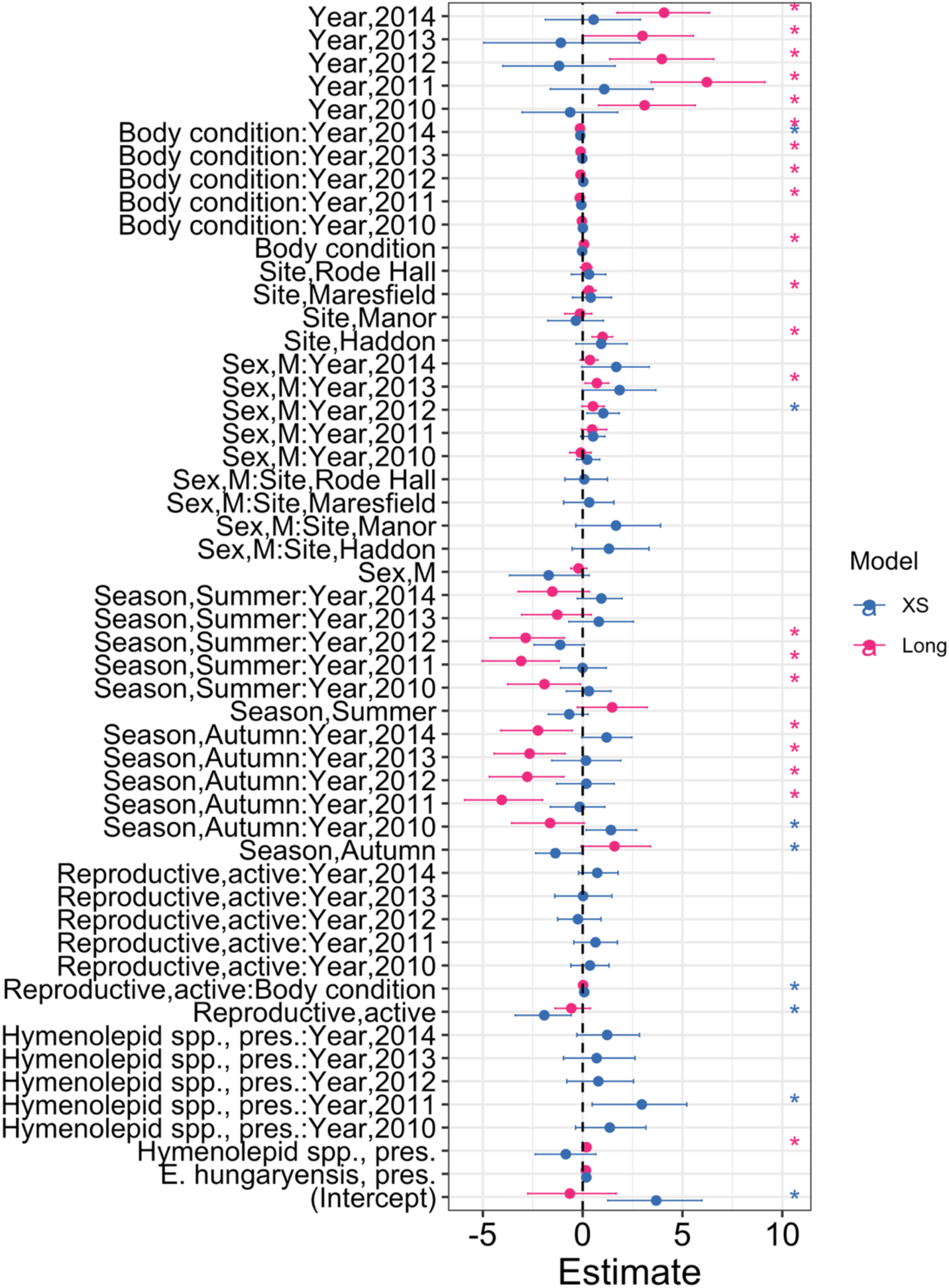
Full model output for optimal model with Interactions remaining in final model after selection. Interactions remaining represent those which had ΔDIC >2. Points and ranges represent model estimates and 95% credibility estimates for each model. Asterisks indicate the significance of variables with a p_MCMC_ <0.05 threshold.

**Figure S7.**
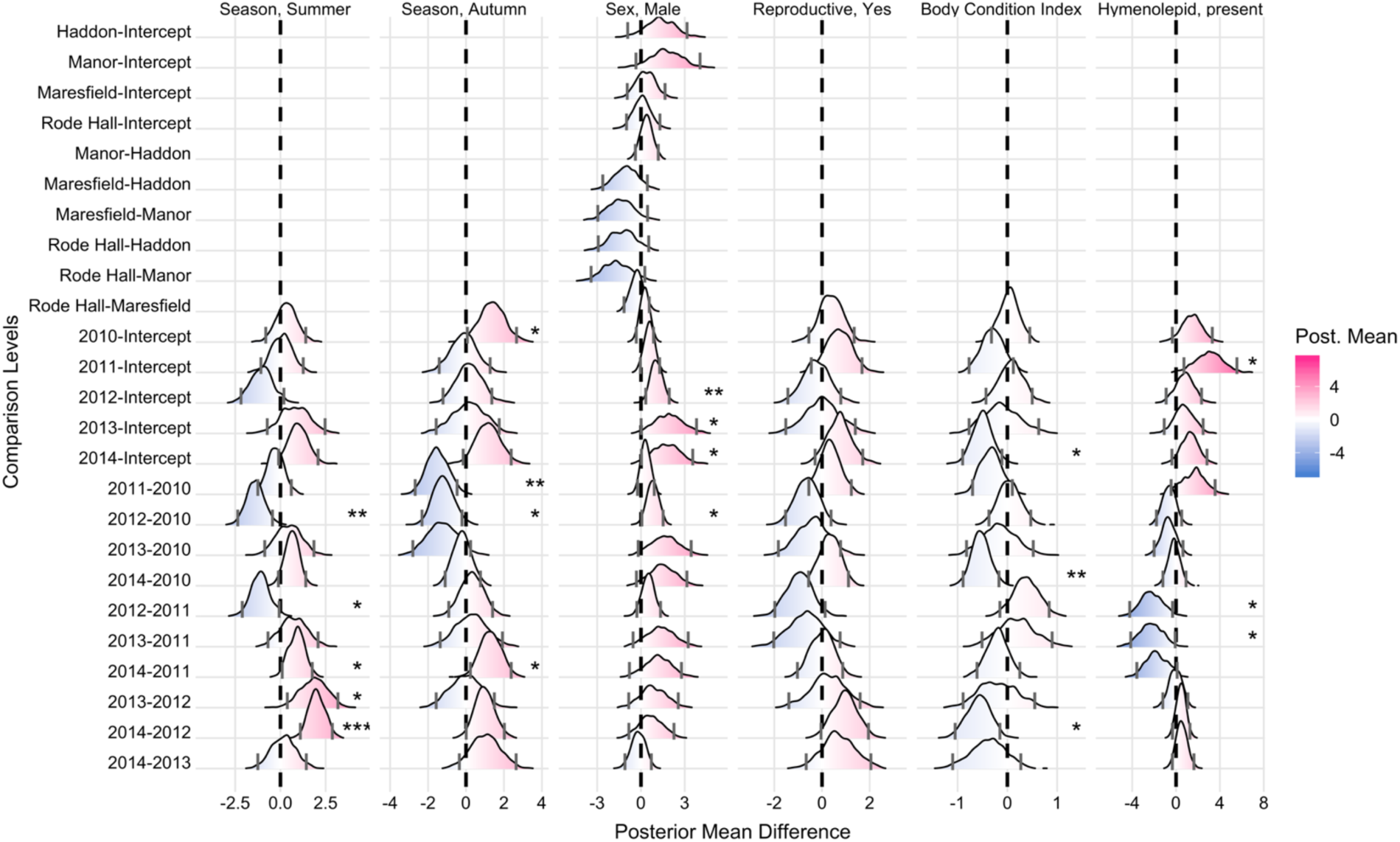
Differences across estimated effects (with 95% credible intervals) for interaction levels which improved full model fit, cross-sectional models. Density ridges represent distributions drawn from the differences between the posterior means of the indicated comparison levels [a-b] for each stored iteration. Blue shading denotes that the slope of effect from the x-axis is lower than that on the y-axis. Differences between effects can be interpreted by comparison of the density ridges to zero; grey lines for each ridge indicate the 95% credibility intervals for these distributions. Blue shading denotes that the slope of effect for [a] is lower than that of [b] for a given interaction. Pink shading denotes that slope of effect from [a] is higher than that of [b]. If credibility intervals do not cross zero, this is considered a significant difference in effect slope of [a-b]. Significant differences between effects are indicates by ***, ** and * for P<0.001, P<0.01 and P<0.05 respectively. ‘Intercept’ represents the baseline year of the model (2009) in all panels.

**Table S4.**
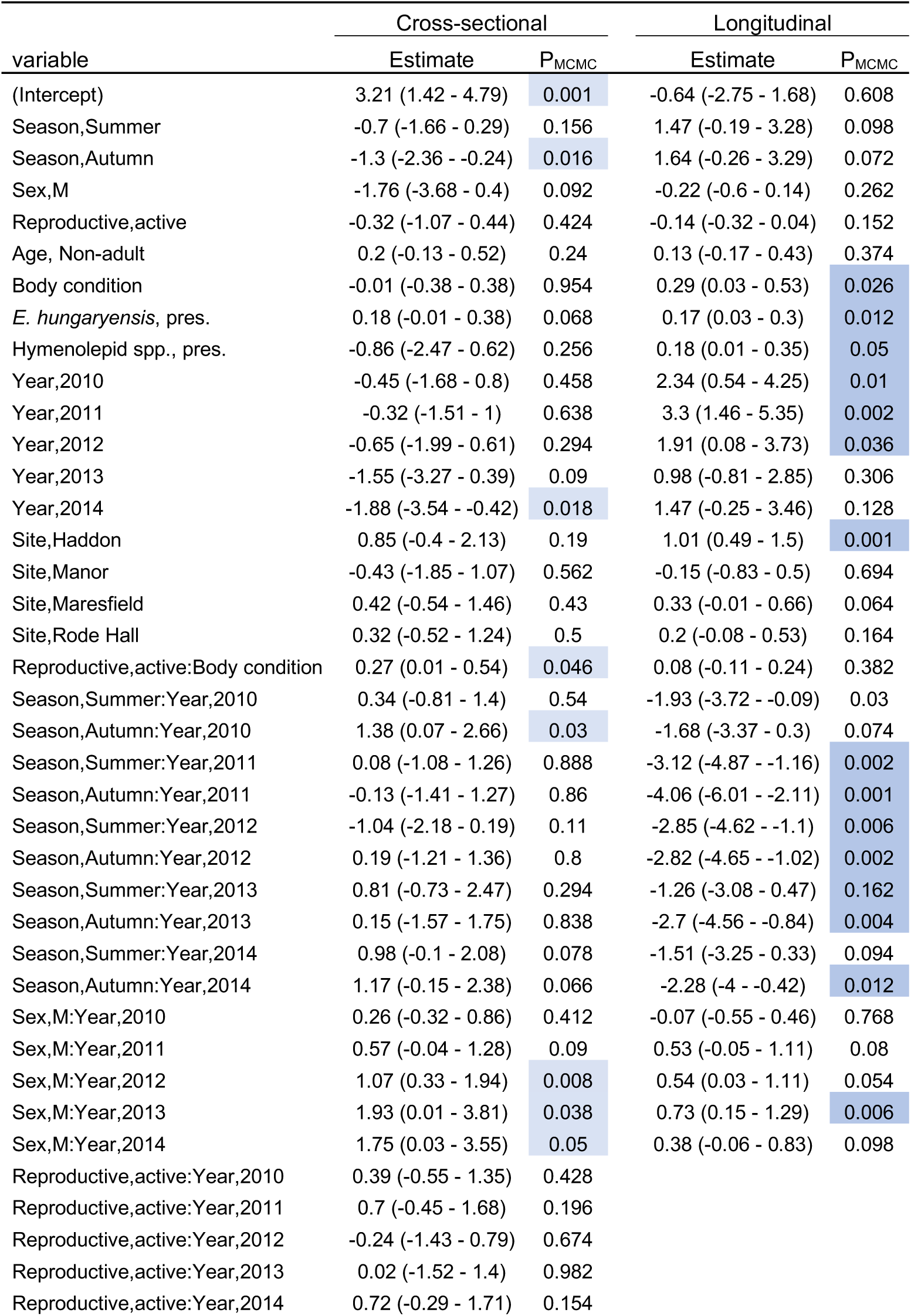
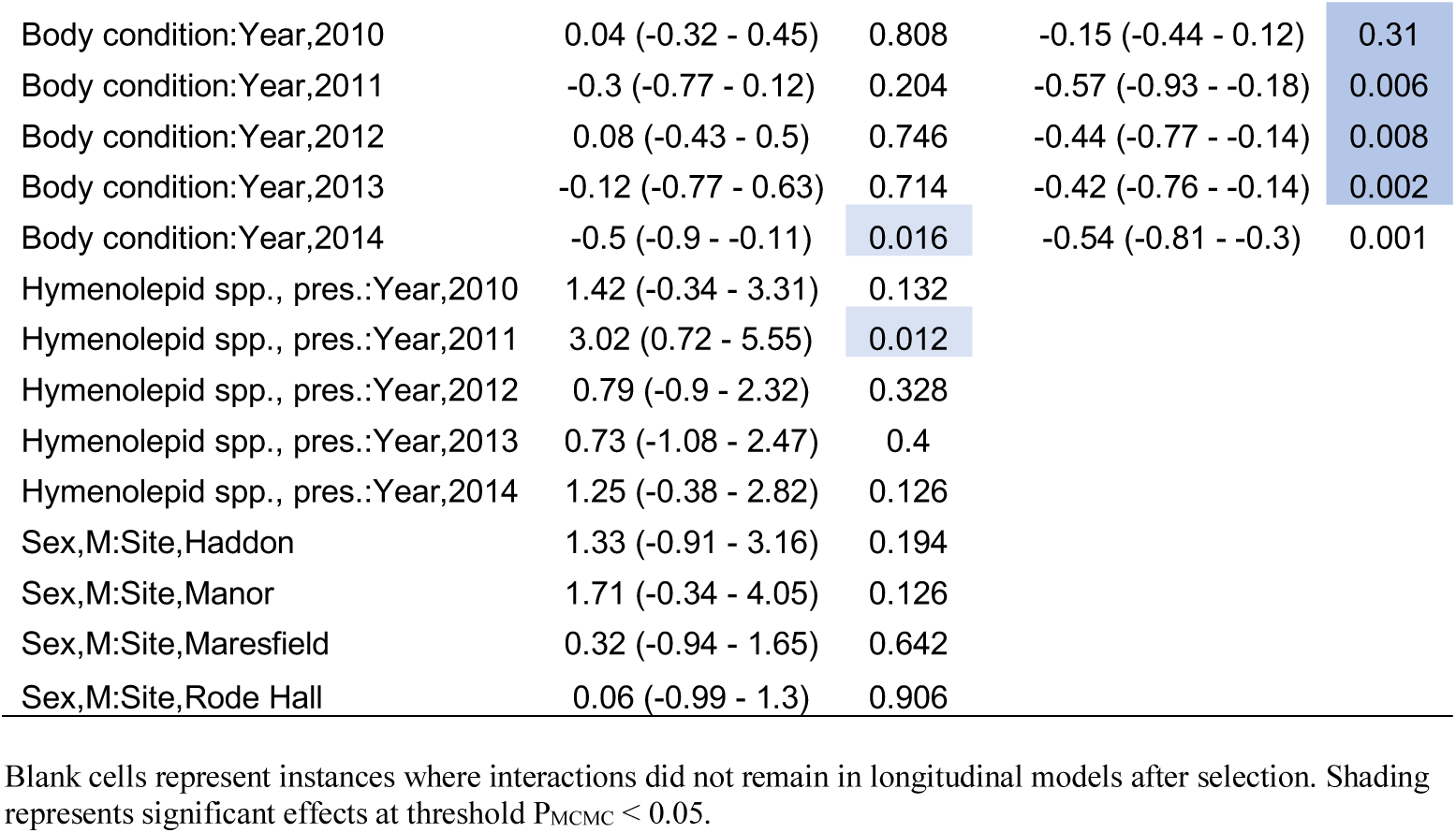
Full Model Output, with interactions.

**Table S5.**
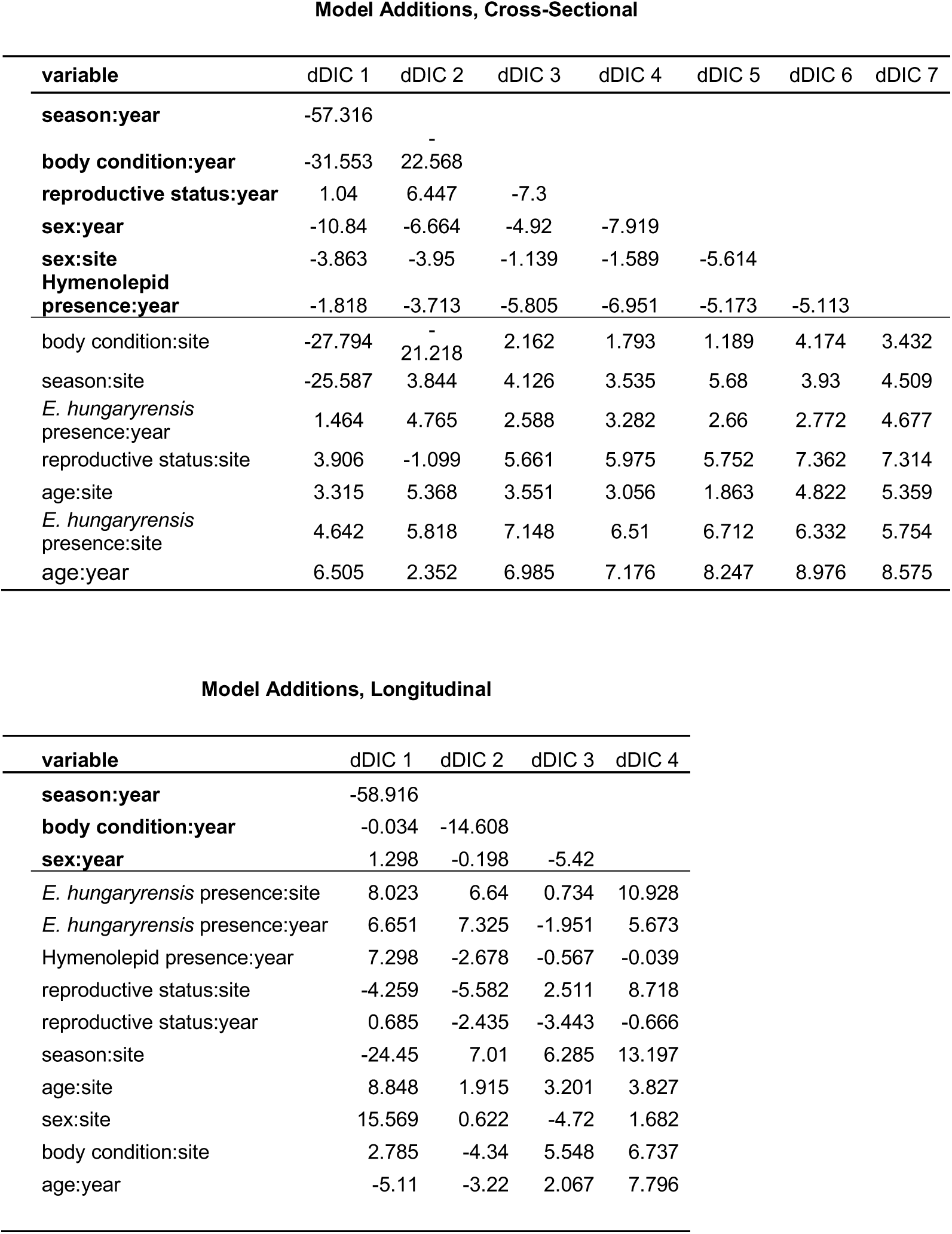
ΔDIC for interactions added to full models for each run of model selection.

**Table S6.**
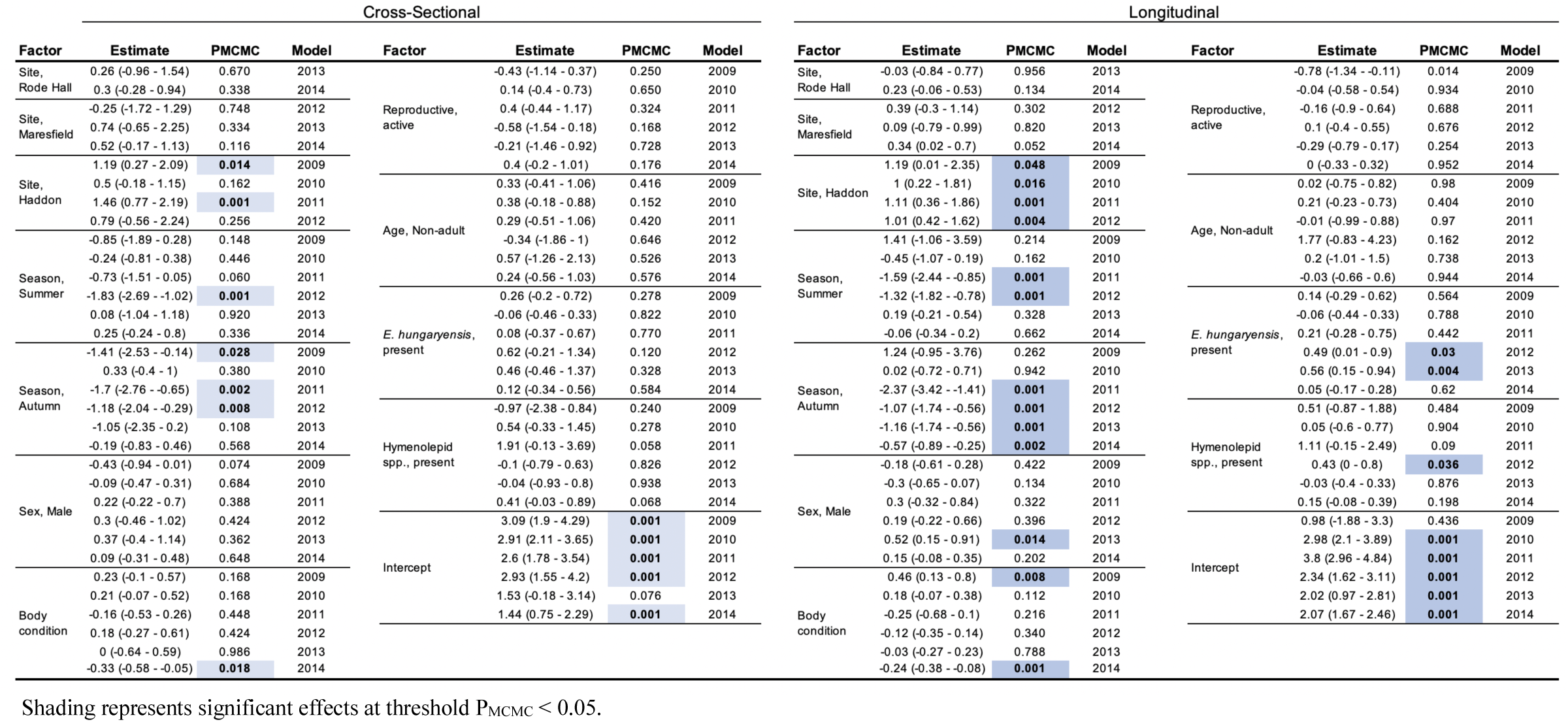
Model output from within-year models.

**Table S7.**
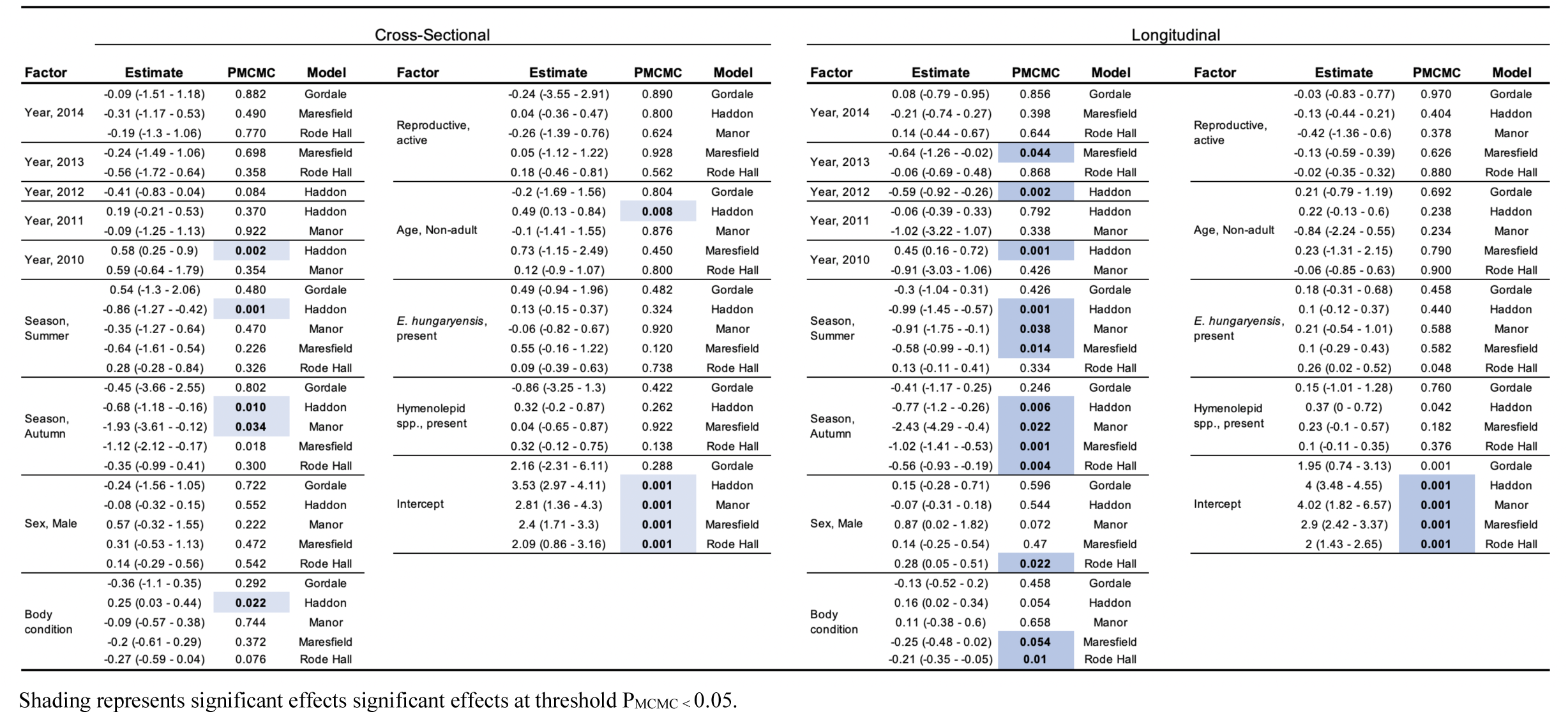
Model output from within-site models.

